## supplementary figures and tables for "Kinetic and structural mechanism for DNA unwinding by a non-hexameric helicase"

**Supplemental Figures**

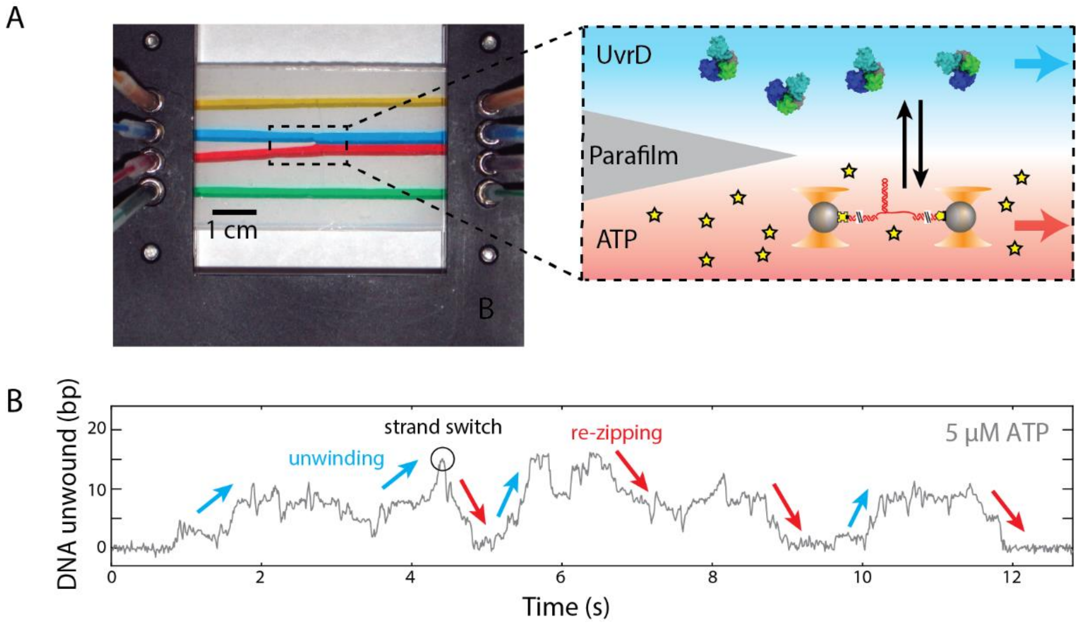


**Supplemental Figure 1. Sample chamber design.** Left: photograph of a laminar flow sample chamber, with different colored food dyes contained in each channel. The polystyrene beads functionalized for tether formation enter the two central channels via thin glass capillaries that connect them to the two outer channels of the chamber (yellow and green in the photograph). Right: Schematic of the two central channels (red and blue in the photograph). Two separate streams containing different buffer components merge at the end of a parafilm taper to form a smooth interface. The top channel contains UvrD but no ATP, while the bottom channel contains ATP but no UvrD. A single DNA hairpin tether is formed in the bottom channel, held at constant force, moved to the top channel to load a UvrD monomer for a 15-30 s incubation period, and finally moved back to the bottom channel to initiate unwinding. Example trace of UvrD unwinding activity at a force of 11 pN and 5 µM ATP. Addition of ATP occurs at ~1 s, leading to UvrD-catalyzed unwinding until dissociation of UvrD at ~12 s. Helicase activity is characterized by short, repetitive cycles of unwinding and re-zipping.

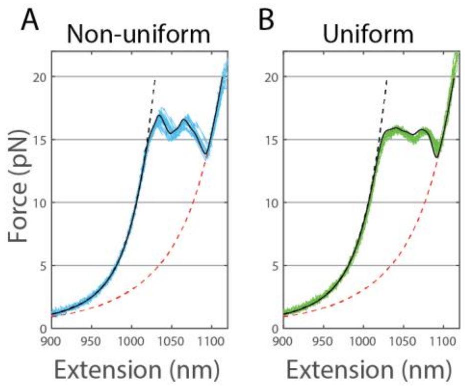


**Supplemental Figure 2. DNA hairpin force-extension behavior.** (**A**) and (**B**) Representative force-extension curves for hairpin sequences with non-uniform (A, light blue) and uniform (B, green) GC distribution in the hairpin stem. Black and red dotted lines represent fits to the extensible worm-like chain model for the closed and open forms of the hairpin, respectively. The solid black line represents a hairpin unfolding model based on the nearest neighbor base-pairing free energies of the stem sequence (see **Materials and Methods**).

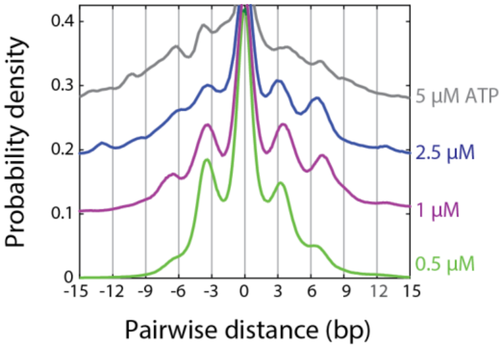


**Supplemental Figure 3. Pairwise distance distributions for unwinding and re-zipping.** Plots are color coded by individual ATP concentration according to the legend. The same intervals from helicase activity traces that were fitted by the step-finding algorithm were compiled to construct the individual pairwise distance plots at each ATP concentration. We use a signed pairwise distance to account for both unwinding and re-zipping. Data is not shown for 10 μM ATP due because high noise and small dwell times make it difficult to discern clear peaks in the pairwise distance distribution at that concentration.

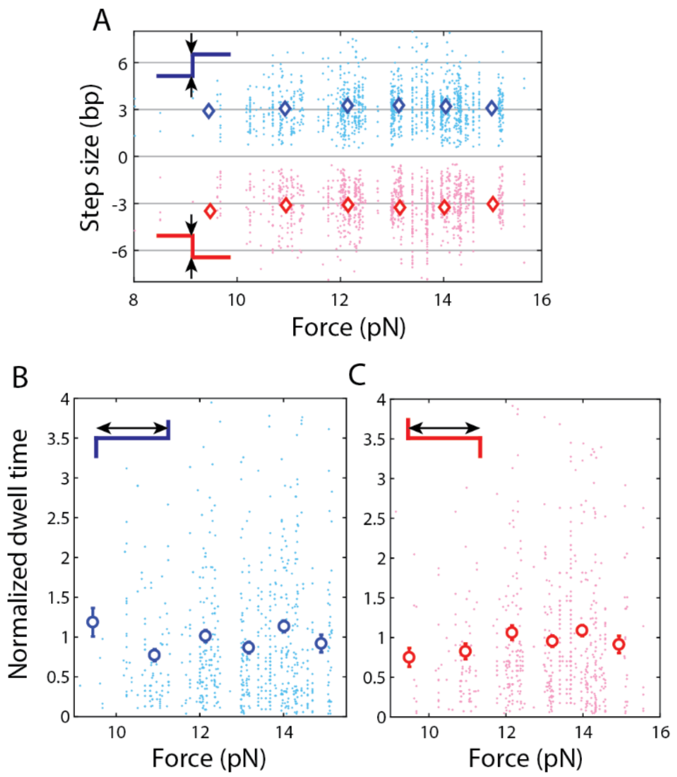


**Supplemental Figure 4. Step sizes and dwell times are independent of applied force. (A)** Scatter plot of individual unwinding (blue dots) and re-zipping (pink dots) step size measurements versus applied force. Averages (colored diamonds) were calculated by binning the data in 1-pN increments across the applied force range, with the exception of data at forces ≤10 pN which were grouped into a single bin due to the smaller number of data points. The s.e.m. is smaller than the diamond symbol size. **(B)** and **(C)** Scatter plots of individual +/+ (blue dots, B) and -/- (pink dots, C) dwell times versus applied force, with averages (colored circles) calculated from the same binning procedure that was used for the step size data. Error bars represent the s.e.m.

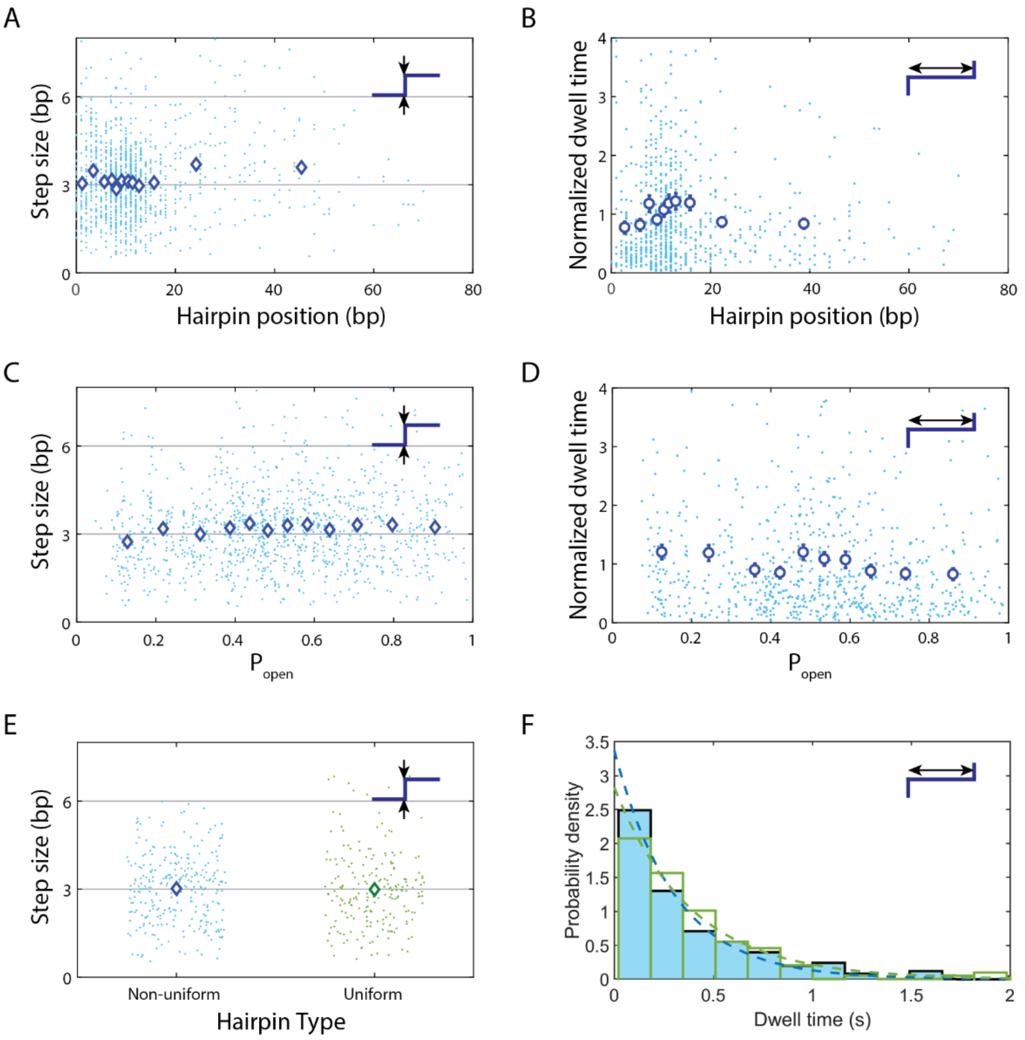


**Supplemental Figure 5. Step sizes and dwell times are independent of DNA sequence. (A)** and **(B)** Scatter plots of individual unwinding step size (blue dots, A) and +/+ dwell time (blue dots, B) measurements versus helicase position along the hairpin sequence. **(C)** and **(D)** Scatter plots of individual unwinding step size (blue dots, C) and +/+ dwell time (blue dots, D) measurements versus $P_{open}$, the thermodynamic probability that one or more base pairs are open downstream of the helicase position along the hairpin stem. Throughout, the plots show boxcar averages over 100 data points for the step size (blue diamonds) and 70 data points for the dwell times (blue circles). **(E)** Comparison of unwinding step size on two hairpin sequences (see **Supplemental Figure 2**). Scatter plot of individual step size measurements for the non-uniform (blue dots) and uniform sequences (green dots) at 1 µM ATP, with averages (colored diamonds). Throughout, error bars denote s.e.m., and the s.e.m. is smaller than the diamond symbol size in the step size plots. **(F)** Comparison of dwell time distributions between the non-uniform (light blue bars) and uniform sequences (green bars) for +/+ step pairs at 1 µM ATP. The dotted lines are fits to a single-exponential function.

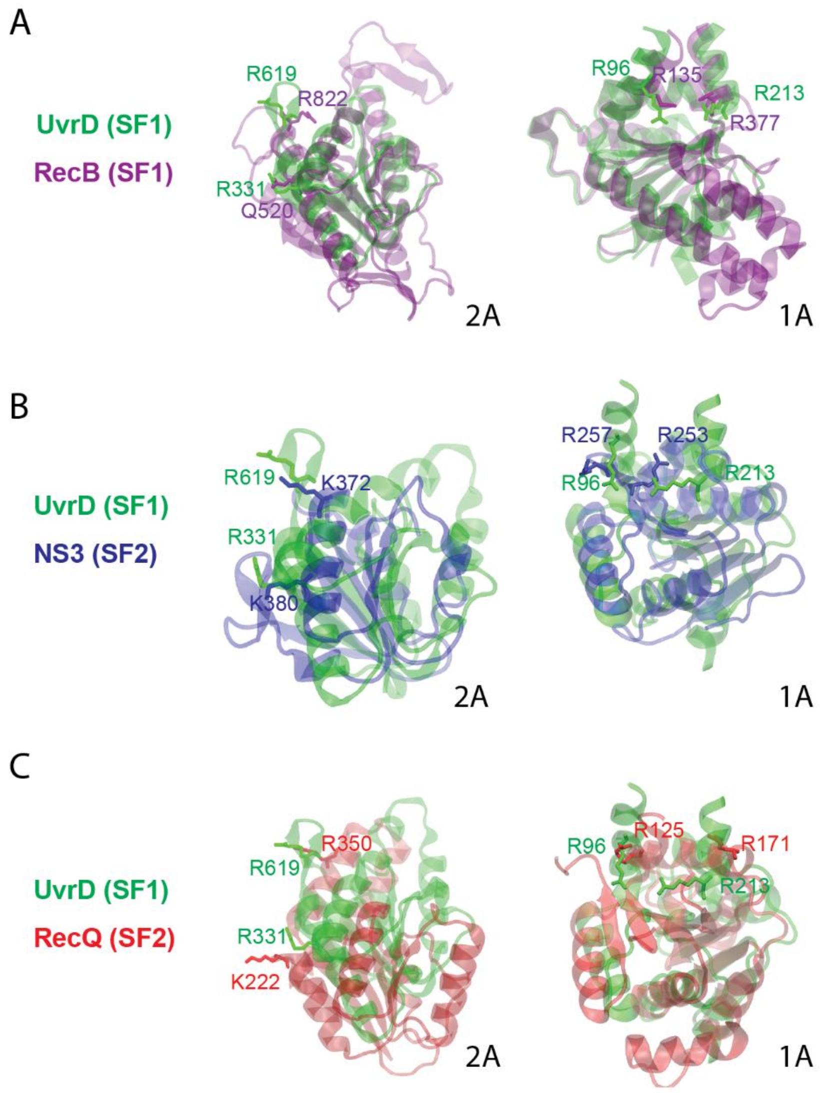


**Supplemental Figure 6. Structural alignment of motor core subdomains of other SF1 and SF2 helicases.** (**A**) Structural alignment of the 1A and 2A subdomains of UvrD (green) and the RecB subunit of SF1 RecBCD helicase (purple). (**B**) Structural alignment of the 1A and 2A subdomains of UvrD (green) and SF2 helicase NS3 (blue). (**C**) Structural alignment of the 1A and 2A subdomains of UvrD (green) and SF2 helicase RecQ (red). Key UvrD residues that participate in loop formation include R619, R331 in the 2A subdomain (left) and R213, R96 in the 1A subdomain (right) and are well preserved in RecB. For NS3 and RecQ, positively charged arginine or lysine residues are found near the locations of those loop-forming residues in UvrD. Residues for each helicase are colored according to the corresponding structure.

**Supplementary Tables**

| **Oligonucleotide function** | **Hairpin identity** | **Sequence (5’-3’)** |
|---|---|---|
| Left handle forward primer | Non-uniform | /5Biosg/ TGA AGT GGT GGC CTA ACT ACG |
| Left handle reverse primer | Non-uniform | CAA GCC TAT GCC TAC AGC AT |
| Right handle forward primer | Non-uniform | 5Phos/ GAC TGA GAC TGA CAT CTC AGA CTG AGA GA TTT TTT TTT T/idSp/ C TCT GAC ACA TGC AGC TCC C |
| Right handle reverse primer | Non-uniform | /5DigN/CAA CAA CGT TGC GCA AAC T |
| Hairpin insert | Non-uniform | 5Phos/CCT GGG GCT GAT AGC TGA GCG GTC GGT ATT TCA AAA GTC AAC GTA CTG ATC ACG CTG GAT CCT AGA GTC AAC GTA CTG ATC ACG CTG GAT CCT ATT TTT AGG ATC CAG CGT GAT CAG TAC GTT GAC TCT AGG ATC CAG CGT GAT CAG TAC GTT GAC TT |
| Left handle forward primer | Uniform | /5Biosg/ TGA AGT GGT GGC CTA ACT ACG |
| Left handle reverse primer | Uniform | CAA GCC TAT GCC TAC AGC AT |
| Right handle forward primer | Uniform | /5Phos/GA CTG TGA CTG ACA TGA GTG ACT GAG ACT TTT TTT TTT T/idSp/CT CTG ACA CAT GCA GCT CCC |
| Right handle reverse primer | Uniform | /5DigN/CAA CAA CGT TGC GCA AAC T |
| Hairpin insert | Uniform | /5Phos/CCT GGA GTC TCA GTC ACT CAT GTC AGT CAC AGT CAG AGT CAT GTC TGA GTC TTG ATG ATG TCA CTG ACT GAG ACT CTG ACT CAC TGA GTC GAG CTT TTG CTC GAC TCA GTG AGT CAG AGT CTC AGT CAG TGA CAT CAT CAA GAC TCA GAC ATG ACT CT |

**Supplementary Table 1. List of sequences for all hairpin inserts and primers.** Abbreviations correspond to the following chemical modifications (readily available from IDT): Phos = phosphate, 5DigN = digoxigenin, Biosg = biotin, idSp = abasic site.

| Sequence | NU | NU | NU | NU | NU | U |
| --- | --- | --- | --- | --- | --- | --- |
| [ATP] (µM) | 0.5 | 1 | 2.5 | 5 | 10 | 1 |
| No. of traces | 25 | 16 | 12 | 12 | 12 | 10 |
| No. of step fitting intervals | 142 | 91 | 65 | 50 | 58 | 60 |
| Total No. of unwinding steps | 394 | 287 | 191 | 134 | 200 | 211 |
| Total No. of re-zipping steps | 338 | 256 | 192 | 138 | 237 | 154 |
| Unwinding step size (bp) | 2.98 ± 0.06 | 3.02 ± 0.07 | 3.33 ± 0.10 | 3.27 ± 0.13 | 3.62 ± 0.11 | 2.99 ± 0.09 |
| Re-zipping step size (bp) | 2.90 ± 0.06 | 3.04 ± 0.08 | 3.11 ± 0.11 | 3.22 ± 0.13 | 3.82 ± 0.13 | 2.99 ± 0.09 |
| Total No. of step pairs | 590 | 452 | 318 | 222 | 379 | 305 |
| Total No. of +/+ dwells | 214 | 156 | 113 | 84 | 153 | 123 |
| Total No. of -/- dwells | 162 | 128 | 105 | 83 | 193 | 71 |
| Mean +/+ dwell time (s) | 0.53 ± 0.04 | 0.37 ± 0.03 | 0.23 ± 0.02 | 0.16 ± 0.02 | 0.08 ± 0.01 | 0.38 ± 0.04 |
| Mean -/- dwell time (s) | 0.42 ± 0.03 | 0.41 ± 0.03 | 0.24 ± 0.03 | 0.16 ± 0.02 | 0.06 ± 0.004 | 0.35 ± 0.04 |

**Supplementary Table 2. Key statistics for step size and dwell time data.** All errors are standard errors of the mean (s.e.m.). Abbreviations: NU = non-uniform sequence; U = uniform sequence.
